# Epigenetic regulation of innate immune memory in microglia

**DOI:** 10.1101/2021.05.30.446351

**Authors:** Xiaoming Zhang, Laura Kracht, Antonio M. Lerario, Marissa L. Dubbelaar, Nieske Brouwer, Evelyn M. Wesseling, Erik W.G.M. Boddeke, Bart J.L. Eggen, Susanne M. Kooistra

## Abstract

Microglia are the tissue-resident macrophages of the CNS. They originate in the yolk sac, colonize the CNS during embryonic development and form a self-sustaining population with limited turnover. A consequence of their relative slow turnover is that microglia can serve as a long-term memory for inflammatory or neurodegenerative events. We characterized the epigenomes and transcriptomes of microglia exposed to different stimuli; an endotoxin challenge (LPS) and genotoxic stress (DNA repair deficiency-induced accelerated aging). Whereas the enrichment of permissive epigenetic marks at enhancer regions explains training (hyperresponsiveness) of primed microglia to LPS challenge, the tolerized response of microglia seems to be regulated by loss of permissive epigenetic marks. Here, we identify that inflammatory stimuli and accelerated aging because of genotoxic stress activate distinct gene networks. These gene networks and associated biological processes are partially overlapping, which is likely driven by specific transcription factor networks, resulting in altered epigenetic signatures and distinct functional (desensitized vs. primed) microglia phenotypes.

## Introduction

Microglia are of myeloid lineage and are long-lived tissue-resident macrophages of the central nervous system (CNS) parenchyma (Prinz et al., 2017).

Macrophages possess innate immune memory (IMM). IMM describes the concept that macrophages, after experiencing a primary ‘priming’ or ‘desensitizing’ stimulus, react with a stronger (immune training) or weaker (immune tolerance) immune response to a subsequent stimulus (Neher & Cunningham, 2019; Netea et al., 2011). IMM was discovered and has been extensively described in blood-derived monocytes/macrophages (Biswas & Lopez-Collazo, 2009; Kleinnijenhuis et al., 2012; Netea et al., 2016; Novakovic et al., 2016; Saeed et al., 2014).

Similar functional states have been described for microglia in mouse models (Eggen et al., 2013; Neher & Cunningham, 2019). Primed microglia can be elicited in mouse models of prion disease (Cunningham et al., 2005), neurodegeneration (Norden et al., 2015; Wendeln et al., 2018), natural aging (Sierra et al., 2007) and neuronal genotoxic stress-induced accelerated aging (D. D. A. Raj et al., 2014). When these mice experienced a peripheral lipopolysaccharide (LPS) challenge, microglia exhibited an excessive immune response manifested by increased expression of proinflammatory cytokines, called microglia training (Cunningham et al., 2005; D. D. A. Raj et al., 2014; Sierra et al., 2007; Wendeln et al., 2018). Oppositely, mouse microglia can be desensitized with LPS (Longhi et al., 2011; Schaafsma et al., 2015; Schaafsma et al., 2017). After a secondary challenge, in the form of LPS (Schaafsma et al., 2015; Schaafsma et al., 2017), traumatic brain injury (Longhi et al., 2011) or cerebral ischemia (J. T. Yu et al., 2010), microglia display immune tolerance which is defined by a reduced immune response.

Microglia are implicated in CNS development, and neurodevelopmental and neurodegenerative diseases (Hammond et al., 2019; Holtman et al., 2015; Kracht et al., 2020; Li et al., 2019; Matcovitch-Natan et al., 2016; Salter & Stevens, 2017; Sarlus & Heneka, 2017). It is especially interesting to investigate microglia IMM in this context. The combination of perturbations like maternal immune activation during vulnerable periods of CNS development together with the occurrence of multiple stimuli over a long period of time is thought to cause neurodevelopmental or neurodegenerative diseases (Knuesel et al., 2014). Microglia are the prime cells that respond to CNS stimuli since they express a wide range of cell surface receptors and adhesion molecules (homeostatic gene signature) through which they can sense those endogenous and exogenous stimuli (Butovsky et al., 2014; Galatro, Holtman, et al., 2017; Gautier et al., 2012; Gosselin et al., 2014, 2017; Hickman et al., 2013; Lavin et al., 2014).

Epidemiologic studies report that infections during specific periods of pregnancy increase the risk for the child to develop neurodevelopmental disorders, like autism or schizophrenia (Estes & McAllister, 2016). Mouse models of maternal immune activation suggest a role for microglia IMM in this process (Knuesel et al., 2014; Neher & Cunningham, 2019). Peripheral LPS challenge of pregnant mouse dams caused preconditioning of offspring microglia which long-lastingly affects microglia LPS responsiveness in adult offspring and also caused behavioral abnormalities (Schaafsma et al., 2017).

In case of neurodegenerative diseases, genetic risk loci are generally immune-related (Cooper-Knock et al., 2017; T. Raj et al., 2014) and specifically enriched in microglia (Nott et al., 2020). A common gene signature was identified in multiple mouse models of neurodegenerative diseases, aging and priming and encompasses genes such as *Axl*, *Clec7a* and *Mac2* (Holtman et al., 2015). This microglia transcriptional phenotype is orchestrated by the APOE-TREM2 pathway and is associated with altered phagocytic and lysosomal activity, and lipid metabolism (Butovsky & Weiner, 2018; Holtman et al., 2015; Keren-Shaul et al., 2017; Krasemann et al., 2017). Given the chronic nature of neurodegenerative diseases, it is hypothesized that microglia are trapped in a primed/trained state ultimately leading to neurotoxicity (Block et al., 2007; Haley et al., 2017). This hypothesis was recently confirmed by the observation that induction of priming of microglia in early adulthood caused exacerbation of Aβ pathology later in life, whereas desensitization of microglia diminished Aβ pathology in a mouse model of AD (Wendeln et al., 2018). Current studies suggest microglia priming and tolerance to have neurotoxic (Schaafsma et al., 2017; Wendeln et al., 2018) or neuroprotective (Longhi et al., 2011; Wendeln et al., 2018; J. T. Yu et al., 2010) consequences, respectively. However, these outcomes should not be generalized and the effects of microglia tolerance and priming on neuronal viability need to be elucidated in a context-specific manner (Neher & Cunningham, 2019).

Both tolerant and trained immunity of peripheral macrophages are long-lasting changes in functionality that are instructed by epigenetic reprogramming (Cheng et al., 2014; El Gazzar et al., 2008; Kleinnijenhuis et al., 2012; Novakovic et al., 2016; Quintin et al., 2012). Though epigenetic programming has been clearly implicated in the segregation of microglia from other tissue resident macrophages in both mouse and human (Gosselin et al., 2014, 2017; Lavin et al., 2014), little is known about the changes in epigenetic signatures in microglia in response to (systemic) immune stimuli or endogenous neuronal damage and how epigenetic memory serves to change subsequent responses. Since microglia are relatively long-lived cells (Füger et al., 2017; Tay et al., 2017), experience of past stimuli is long-lastingly secured in the microglial epigenome and can thus have persistent consequences on microglia functionality and neuronal viability. Several lines of evidence suggest a role for epigenetic regulation of microglia functional states (Cho et al., 2015; Keren-Shaul et al., 2017; Matcovitch-Natan et al., 2016; Matt et al., 2016; Schaafsma et al., 2015; Wendeln et al., 2018; Yeh & Ikezu, 2019).

To delineate the gene networks and associated epigenetic signatures and transcription factors that underlie functional microglia states of priming and tolerance, we acutely isolated microglia from mice challenged with LPS and from accelerated aging mice and analyzed their transcriptional and chromatin status at a genome-wide level.

## Results

### LPS desensitization and accelerated aging result in distinct transcriptional responses in microglia

In mouse, previous data indicated two distinct microglia functional states of ‘desensitization’ induced by an intraperitoneal LPS challenge (Schaafsma et al., 2015) and ‘priming’ during accelerated aging resulting from deficiency of the DNA-damage repair protein Ercc1 (D. D. A. Raj et al., 2014). These different functional states can be unmasked by a (secondary) LPS stimulus resulting in a ‘tolerant’ or ‘trained’ immune response, respectively, and were so far characterized based on the analysis of limited sets of genes by qPCR (D. D. A. Raj et al., 2014; Schaafsma et al., 2015). For several tested inflammatory genes, such as *Il1b*, *Tnf*, and *Il6*, the initial stimulus determined whether microglia show a dampened or enhanced response to (secondary) LPS treatment. However, the genome-wide transcriptional remodeling in desensitized and primed microglia and its effect on responsiveness to future inflammatory exposure are unknown. Therefore, we performed RNA-sequencing on acutely isolated, FACS-purified microglia from mice that were either recurrently treated with LPS with a 1-month interval, or from *Ercc1^Δ/ko^* mice that were stimulated with LPS near the end of their lifespan at 10-12 weeks of age (Figures 1, S1, S2A, S2B, Supplemental file 1, 2).

**Figure 1.**
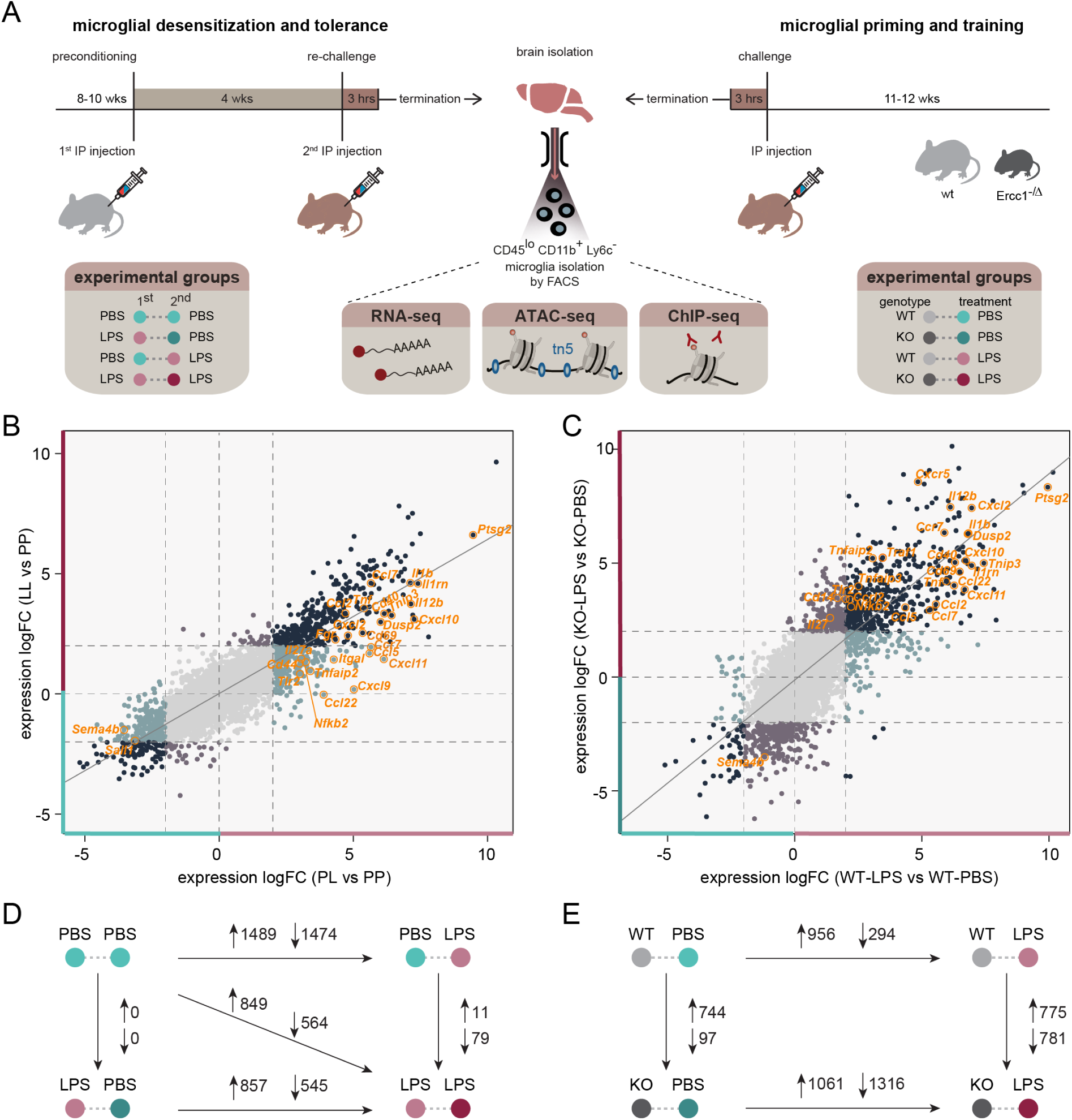
LPS desensitization and accelerated aging result in distinct changes of the microglia immune response. **a,** Graphic representation of the mouse models and treatment groups. A pure microglia population was isolated by FACS and subjected to RNA, ATAC, and ChIP-sequencing analysis. **b, c,** Four-way plots depicting changes in gene expression in microglia isolated from LPS-injected naive and pre-conditioned mice (n=3 per experimental condition) (**b**) and *Ercc1^Δ/ko^* and control mice (n=3 per experimental condition) (**c**). Every gene is represented by an individual dot. Differentially expressed genes (LogFC>2) are labeled with different colors indicating their respective expression changes. Dark blue dots indicate genes differentially expressed in both comparisons; turquoise (PL versus PP and WT-LPS versus WT-PBS) and lavender (LL versus PP and KO-LPS versus KO-PBS) dots represent genes differentially expressed in one of the comparisons, several relevant genes are highlighted. **d, e,** The number of differentially expressed genes (LogFC>1 and FDR<0.01) between treatment groups in the endotoxin tolerance (**d**) and *Ercc1^Δ/ko^*-induced microglia priming models (**e**). Upward arrows indicate increased gene expression, downward arrows indicate decreased gene expression in the condition where the large arrow points to.

For the tolerance model, we analyzed four treatment groups: the controls that were treated with PBS twice (PP), mice that were treated with LPS and after 1 month with PBS (LP) to investigate desensitization, mice treated with PBS followed by LPS after 1 month to determine the acute response to LPS (PL) and mice that were treated with LPS twice with a 1-month interval between challenges (LL) to identify the tolerant response (Figure 1). I.p. injection of LPS resulted in significant changes in gene expression in microglia after 3 hours (Figures 1B, 1D, S2A). After 1 month, this initial response to LPS had subsided and in terms of the transcriptional program, no significant differences were observed between the PP and LP groups (Figures 1D, S2A). However, when mice were challenged with LPS for a second time, the response was different from the initial response (Figure S2A) and many genes were significantly differentially expressed between PL and LL conditions (Figure 1B, 1D).

For the microglia priming model, both the *Ercc1^Δ/ko^* mice and their control littermates were treated with PBS (WT-PBS, KO-PBS) to identify priming effects or with LPS (WT-LPS, KO-LPS) to identify training. As we observed previously, deletion of *Ercc1* in itself results in significant changes in gene expression (Figures 1E, S2B). However, when *Ercc1^Δ/ko^* mice were treated with LPS, the difference between microglia from control and knockout mice was much more pronounced (Figures 1C, 1E, S2B).

The response to an acute LPS stimulus was highly similar in mice of the tolerance (C57BL/6J) and priming (FVB/ C57BL/6J) model. Nevertheless, it cannot be fully excluded that the LPS response is slightly affected by the genetic backgrounds of the two mouse strains used in this study.

Many genes differentially expressed between PP versus LP and WT-PBS versus WT-LPS showed very similar changes in expression in response to LPS, after ranking them based on expression level and comparing rank positions between the two groups (Figure S2C). With our RNA-sequencing dataset, we confirmed several of our previous findings (Holtman et al., 2015; D. D. A. Raj et al., 2014), and replenish this information with complete gene expression profiles of the desensitized and primed microglia phenotype. The opposite regulation of the pro-inflammatory genes *Il1b* in tolerant (LL) and trained (KO-LPS) microglia (Figure S2D, S2E) was confirmed. In addition, primed microglia (KO-PBS) showed increased expression of genes belonging to the ‘primed’ gene hub (Holtman et al., 2015), including *Clec7a* and *Axl* when compared to control animals (WT-PBS, Figure S2E).

### Genes with distinct transcriptional responses to LPS have different biological functions

Following 3 hours of LPS exposure (PL versus PP, LogFC>1, FDR <0.01), 1489 genes showed increased expression (Figures 1D, 2A) while 1474 genes were downregulated in microglia (Figures 1D, S3A). Generally, LPS-induced genes were involved in various aspects of the immune response (Figure S3B, Supplemental file 3), while genes downregulated by LPS were involved in multiple biological processes (Figure S3C, Supplemental file 3). Of note, in the LPS-downregulated genes, the association with biological processes showed a lower level of significance than the GO terms associated with LPS-upregulated genes (Figures S3B, S3C, Supplemental file 3).

**Figure 2.**
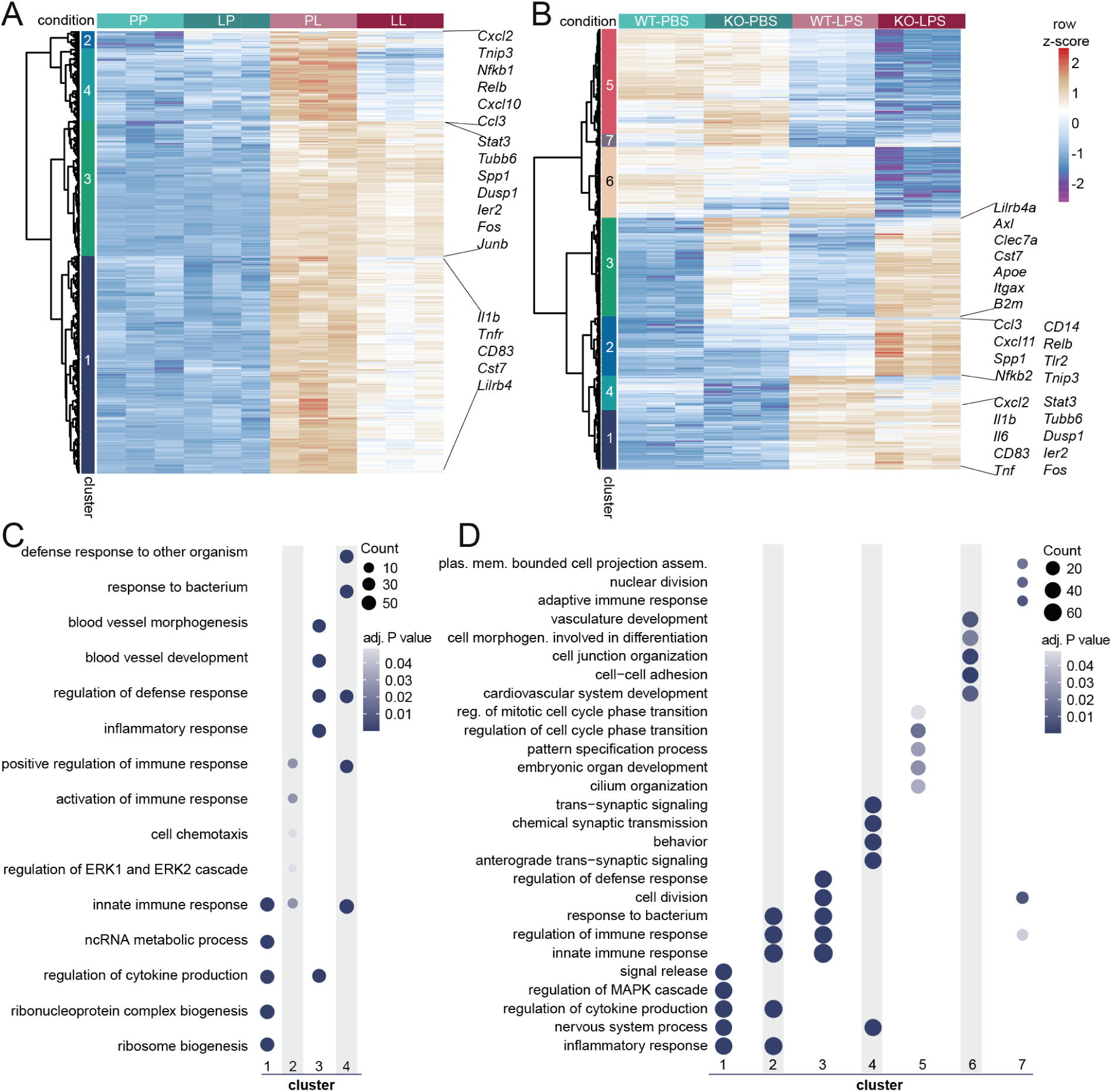
Identification of gene clusters with distinct transcriptional programs in desensitized and primed microglia. **a, b,** Heatmaps with Manhattan distance-based hierarchical clustering analysis of upregulated genes in response to LPS in microglia of C57/BL6 mice three hours after i.p. injection with LPS (LogFC>1 and FDR<0.01, PL versus PP). Four main clusters are identified, containing tolerized genes (cluster 2 and 4) and responsive genes (cluster 1 and 3) that show distinct activity to LPS re-stimulation. **b,** Heatmap with Manhattan distance-based hierarchical clustering analysis of all genes differentially expressed between *Ercc1^Δ/ko^* (KO) and *Ercc1^wt/ko^*, *Ercc1^wt/Δ^, Ercc1^wt/wt^* (WT) mice with or without LPS injection (n=3 per experimental condition). Seven clusters are identified, including two clusters of genes primed and trained to LPS treatment in KO mice (cluster 3 and 2). **c, d,** Top 5 GO annotations, based on gene count per GO term, of responsive (cluster 1, 3) and tolerized (cluster 2, 4) gene clusters (**c**) and the 7 clusters identified in *Ercc1^Δ/ko^* microglia (**d**).

Focusing on LPS-induced genes, out of 1489 genes, 1187 responded similarly in case of re-stimulation with LPS (cluster 1 and 3), while 302 showed a reduced response to a second LPS challenge (cluster 2 and 4, Figure 2A). Processes uniquely associated with the 1187 responsive genes were ‘ribosome biogenesis’, ‘regulation of cytokine production’, and ‘inflammatory response’, while ‘positive regulation of immune response’ and ‘response to bacterium’ were particularly associated with the 302 tolerized genes (Figure 2C, Supplemental file 3). This is in line with the finding that tolerant monocytes/macrophages are impaired in their ability to produce pro-inflammatory cytokines (Foster et al., 2007; Novakovic et al., 2016), but are capable of expressing genes involved in damaging or killing pathogens, so-called antimicrobial effectors. These data suggest that an i.p. injection with LPS initially induces a major immune response in microglia, which then results in the establishment of long-term innate immune tolerance that is characterized by a significantly reduced transcriptional response to secondary LPS treatment.

### Primed microglia have a genome-wide exaggerated response to LPS treatment

To gain insight into the biological processes affected by *Ercc1* deletion in microglia from unstimulated and LPS-treated mice, Manhattan distance-based hierarchical clustering analysis of genes followed by gene ontology analysis per cluster was performed (Figures 2B, 2D, Supplemental file 4). Seven clusters were identified containing genes that were altered by Ercc1 deletion (KO). Genes of clusters 5 are similarly affected in WT and KO microglia and downregulated in both genotypes after LPS treatment. GO terms associated with these genes included ‘regulation of cell cycle phase transition’, and ‘pattern specification process’. The expression of genes in cluster 7 are induced in KO compared to WT microglia and are depleted in both conditions after LPS treatment. These genes are involved in processes like ‘nuclear division’, ‘cell division’ and ‘the immune response’. Cluster 6 contain genes that are upregulated in microglia of PBS and LPS-treated WT compared to KO mice. These genes are associated with ‘cell junction organization’ and ‘cell-cell adhesion’.

Cluster 3 contains genes that were induced in PBS-treated and to a greater extent in LPS-treated KO compared to WT microglia. These primed genes are associated with GO terms ‘regulation of defense response’, ‘cell division’, ‘response to bacterium’ and ‘innate immune response’. Cluster 2 contains genes that were induced by LPS in KO and to a lesser extent in WT microglia and these genes were associated with GO terms such as ‘response to bacterium’, ‘innate immune response’ and ‘regulation of cytokine production’, underlining the trained immune response of KO microglia to LPS challenge. Cluster 4 contains genes that were induced by LPS in WT and to a lesser extent in KO microglia and these genes were associated with GO terms such as ‘trans-synaptic signaling, ‘chemical synaptic transmission’ and ‘nervous system process’. Finally, genes in cluster 1 are induced by LPS to a similar degree in WT and KO microglia and are associated with GO terms, like ‘signal release’, ‘regulation of cytokine production’ and ‘inflammatory response’ (Figure 2D).

In agreement with our previous findings (D. D. A. Raj et al., 2014), also at a genome-wide level, Ercc1 deficiency generates an environment where microglia are more responsive to inflammatory stimuli, as evidenced by a large set of inflammatory genes whose expression is significantly increased in microglia upon LPS treatment of *Ercc1^Δ/ko^* mice.

### Epigenetic remodeling in response to LPS desensitization and accelerated aging

The transcriptomes of microglia from PP and LP treated mice are almost identical, however, they respond very differently to re-stimulation with LPS (Figures 2A, S2A). Similarly, many genes that are not transcriptionally altered in Ercc1 deficient mice show an increased transcriptional response to LPS (Figures 2B, S2B). These data suggest that microglia have innate immune memory that is not secured in their transcriptome. Rather, similar to macrophages and as suggested by our previous analysis of the *Il1β* locus (Schaafsma et al., 2015), it is likely that epigenetic reprogramming is involved.

To gain insight in the genome-wide epigenetic changes induced by LPS desensitization and Ercc1 deficiency, we performed assay for transposase accessible chromatin-sequencing (ATAC-seq), which indiscriminately identifies open chromatin regions in the genome (Buenrostro et al., 2013, 2015), and chromatin immunoprecipitation-sequencing (ChIP-seq), which probes histones carrying specific posttranslational modifications (Henikoff & Shilatifard, 2011; Kouzarides, 2007). In case of the tolerance model, we used antibodies targeting H3K4me3 and H3K27Ac to identify transcription start sites (TSSs) and enhancers of actively transcribed genes, respectively. In *Ercc1^Δ/ko^* mice, we also analyzed H3K4me3 and H3K27ac, and additionally H3K4me1 which together with H3K27ac marks active enhancers and the Polycomb-regulated H3K27me3 associated with transcriptional repression (Figure S4A).

Representative examples of chromatin accessibility and occupation, and RNA expression of individual tolerized (*Il1b*, *Tnf*, *Ccl3, Nfkb1* and *Relb,* Figure S2D, S4B) and primed/trained (*Il1b, Ccl3, Cxcl11*, *Clec7a* and *Axl,* Figure S2E, S4C) genes are depicted and indicate dynamic regulation of epigenetic signatures associated with changes in gene expression levels.

### Epigenetic characterization of tolerized genes

In order to determine which chromatin characteristics correspond to the transcriptional changes induced by LPS, we identified regions in the genome with significant differences in chromatin accessibility or histone modifications. Differential peaks were classified as promoters when they were located within 1000 bp downstream and 1000 bp upstream of a TSS of the nearest gene and as enhancers when being located distal of this region. To integrate RNA-, ATAC-, and ChIP-seq data, the differentially expressed genes (logFC) were correlated to differentially regulated chromatin regions (M-value) within one comparison.

Similar to what has been described in macrophages (Escoubet-Lozach et al., 2011; Hargreaves et al., 2009; Saeed et al., 2014), in microglia H3K4me3 already marks TLR4-responsive promoters prior to LPS stimulation (Figure S4B). Irrespective whether microglia are exposed to LPS for the first or the second time, genes which are expressed in response to LPS are, except for a small group of tolerized genes, largely overlapping (Figure 2A). LPS-induced gene expression significantly correlates with ATAC, H3K4me3, H3K27Ac peak enrichment, associated with permissive gene expression, in promotors and enhancers of microglia from PL compared to PP (Figure 3A left panel, Supplemental file 5) and LL compared to LP-treated mice (Figure 3A middle panel, Supplemental file 5). In line with the fact that H3K4me3 is generally associated with promoters, LPS-induced gene expression only significantly correlates with enrichment of this mark in promoters but not enhancers in the PL versus PP comparison (Figure 3A left panel, Supplemental file 5).

**Figure 3.**
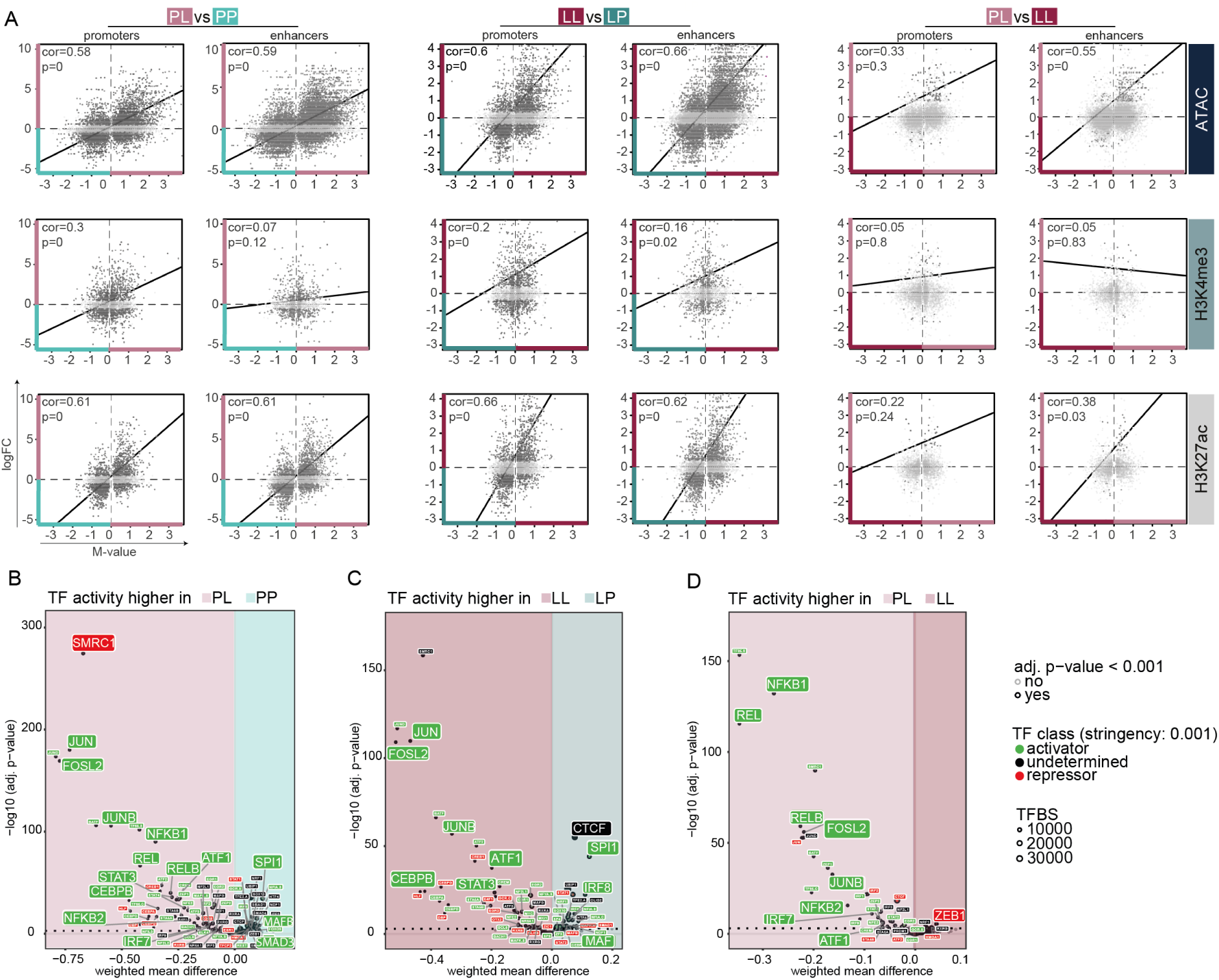
The LPS response in naive and desensitized microglia is defined by enhancer signatures of transcriptional permissive marks. **a,** Scatterplots depicting the correlation of differentially expressed genes (logFC) with corresponding differential ATAC, H3K4me3 or H3K27ac peaks (M-value) at promoters (within 2 kb of the nearest TSS) or enhancers (distal to promoters) between PL versus PP (left panel), LL versus LP (middle panel), PL versus LL (right panel). Each dot represents a differentially expressed gene that is associated with a significant differential chromatin peak (FDR<0.0)1 in the given comparison. Light gray-colored dots indicate non-significant gene expression differences (FDR>0.01). **b, c, d,** Transcription factor binding site analysis generated by diffTF to identify critical regulators for different gene sets based on ATAC- and RNA-seq data. Volcano plots depicting weighted mean difference of accessible TFBS between PL versus PP (**b**), LL versus LP (**c**), or PL versus LL (**d**). The color of each TF indicates its classification into an activator (green), a repressor (red) or undetermined (black) based on correlation of TFBS accessibility with RNA expression of the TF. FC=fold change, TF=transcription factor, TFBS=transcription factor binding site.

Tolerized genes are characterized by increased expression after the primary LPS challenge (PL) and reduced induction after the secondary LPS challenge (LL) (Figure 2A). When comparing the microglia response to primary and secondary LPS challenge (PL versus LL), the expression of the tolerant genes after primary LPS challenge significantly correlated with enrichment of ATAC and H3K27ac peaks at enhancers, but not promoters (Figure 3A right panel, Supplemental file 5). This means vice versa that after secondary LPS challenge, tolerized genes were depleted in these activating expression-associated enhancer marks. The expression of tolerized genes was not significantly correlated to the promoter-associated histone mark H3K4me3. Together, these results indicate that the tolerized response of microglia to LPS seems to be mainly enhancer driven and, at least partially, explained by a loss of histone marks associated with active expression after secondary LPS challenge (Figure 3 right panel, Supplemental file 5).

Transcription factors (TFs) are critical determinants of changes in both transcriptional and epigenetic programs that can be activated by signaling pathways. TFs are often part of large, multimeric protein complexes that also contain chromatin-modifying enzymes, and recruitment of TFs can result in local remodeling of the chromatin (Zhou et al., 2017). DiffTF was used to identify the TFs that might be involved in the differential chromatin regulation in tolerant microglia. Differential chromatin accessibility peaks (weighted mean difference) of putative TF binding sites (TFBS) between two conditions were identified. Next, this ATAC-seq data was integrated with RNA-seq data by correlating differential accessible peaks of putative TFBS to differential gene expression levels of a particular TF. This procedure is then repeated for each TF. Based on whether the correlation of TF activity and expression is positive or negative, TFs were classified as an activator or a repressor. Alternatively, when there was no correlation detected, the TF was classified as undetermined or the TF was not expressed (Berest et al., 2019, Figure 3B, 3C, 3D).

Genome-wide accessible chromatin regions, significantly enriched in naïve microglia (PP, Figure 3B, Supplemental file 6), contain binding sites for the key myeloid TFs PU.1 (SPI1), IRF8 and MAFB, described to be crucial for adult mouse microglia transcriptional identity (Kierdorf et al., 2013; Matcovitch-Natan et al., 2016). In addition, naïve microglia are enriched in TFBS for SMAD3, an effector molecule downstream of TGFβ (Massagué et al., 2005), which is critical for the microglia homeostatic signature (Butovsky et al., 2014). Binding sites of homeostasis-associated TFs were lost and TFBS of known mediators of LPS-induced inflammatory pathways in macrophages/microglia (Escoubet-Lozach et al., 2011; Holtman et al., 2015; Roach et al., 2007) including the NF-κB TF family (NFKB1/2, REL/RELB, Oeckinghaus & Ghosh, 2009), TFs involved in the immediate early response (IER; JUN, JUNB, FOSL2) and the interferon pathway (IRF TF family), STAT3, CEBPB (Simpson-Abelson et al., 2017; Straccia et al., 2011; Valente et al., 2013), and the general activating transcription factor ATF1 were all detected to be enriched in microglia acutely challenged with LPS (PL versus PP, Figure 3B, Supplemental file 6).

After a secondary LPS challenge, in tolerant microglia (LL vs. LP), many of these inflammatory-associated TFBS are still enriched, except those belonging to the NF-κB TF family, indicating that recruitment of these TFs specifically occurs after primary LPS challenge. This is also confirmed in the direct comparison of acutely stimulated versus tolerant microglia (PL vs. LL, Figure 3D, Supplemental file 6). Furthermore, the enrichment of TFBS for SPI1, IRF8, CTCF and MAF, important for the homeostatic microglia transcriptome (Gosselin et al., 2014; Matcovitch-Natan et al., 2016), in desensitized microglia (LP vs. LL, Figure 3C, Supplemental file 6) explains their naive-like transcriptome (Figure 1D).

Many of the inflammatory-associated putative TFBS are depleted and the TFBS for the transcriptional repressor ZEB1, associated with suppression of immune active genes (C. J. Block et al., 2019; Wang et al., 2009), are enriched in tolerant microglia (LL) when compared to microglia of acutely LPS-challenged mice (PL, Figure 3D, Supplemental file 6), possibly explaining the dampened expression of tolerized genes in LL microglia.

These data indicate that deposition of permissive chromatin marks drive the acute LPS-response of microglia, and loss of those, in particular surrounding TFBS of the NF-κB family, might at least partially explain the tolerized response of microglia to a secondary LPS-challenge.

### Epigenetic characterization of the priming response

In case of microglia priming, we also observed a general concordance between the transcriptional changes following Ercc1 deficiency and LPS challenge and the presence of permissive chromatin characteristics. Induction of gene expression by *Ercc1* KO or by LPS in both WT and KO microglia significantly correlated with increased chromatin accessibility in promoters as well as enhancers (Figure 4A, Supplemental file 2 and 7). In addition, compared to WT-PBS, many KO-induced genes are marked with significant enrichment of the permissive marks H3K27Ac and H3K4me3 (Figure 4B). The expression of some of the KO-induced genes additionally correlates with H3K4me1 enrichment, which together with H3K27ac deposition is associated with active transcription (Creyghton et al., 2010). Inversely, some of the genes whose expression is induced by Ercc1 deficiency are depleted in H3K27me3, which is associated with Polycomb-associated gene repression, at promoters of microglia from KO versus WT mice (Figure 4B). Together, this indicates that the expression of primed genes in Ercc1 deficient mice is driven by enriched chromatin characteristics associated with permissive transcription and depletion of repressive chromatin marks.

**Figure 4.**
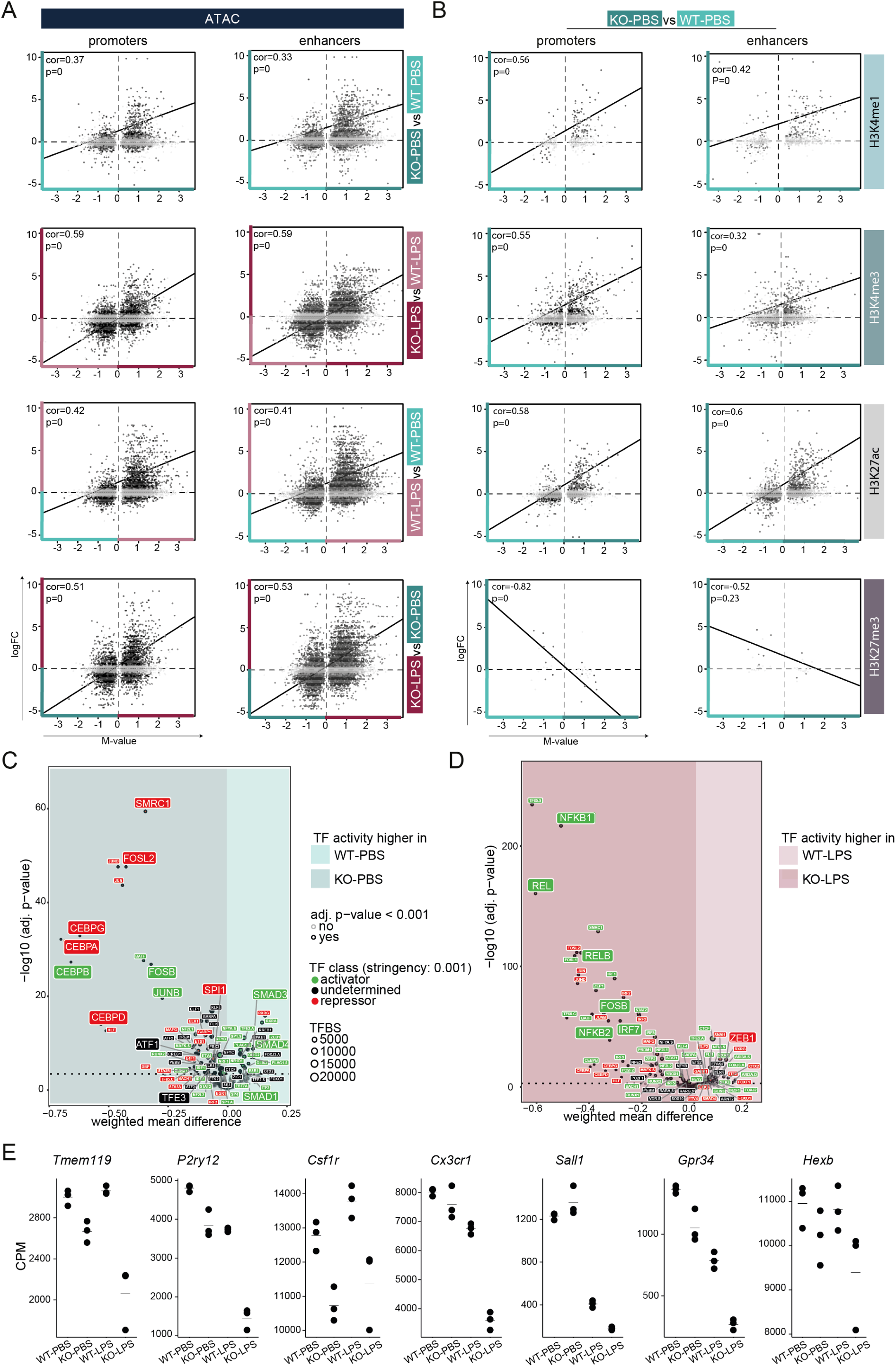
Enhancer and promoter signatures of transcriptional permissive marks regulate training in primed microglia. **a, b,** Scatterplots depicting the correlation of differentially expressed genes (logFC) with corresponding differential ATAC peaks (M-value) in KO versus WT, LPS-treated KO versus LPS-treated WT, LPS-treated WT versus WT and LPS-treated KO versus KO microglia (**a**), and differential H3K4me1, H3K4me3, H3K27ac or H3K27me3 peaks (M-value) in KO versus WT microglia (**b**). The chromatin peaks are divided into promoters (blue, within 2 kb of the nearest TSS) and enhancers (purple, distal to promoters). Each dot represents a differentially expressed gene that is associated with a significantly differential chromatin peak (FDR<0.01) in the given comparison. Gray color of dots indicates non-significant gene expression differences (logFC >1, FDR>0.01). **c, d,** Transcription factor binding site analysis generated by diffTF to identify critical regulators for different gene sets based on ATAC- and RNA-seq data. Volcano plots depicting weighted mean difference of accessible TFBS between KO-PBS versus WT-PBS (**c**) and KO-LPS versus WT-LPS (**d**) microglia. The color of each TF indicates its classification into activator (green), repressor (red) or undetermined (black) based on correlation of TFBS accessibility with RNA expression of the TF. **e,** Gene expression values (CPM, Supplemental file 2) of selected homeostatic microglia genes in the primed mouse model. Every dot depicts an individual animal (n=3 per experimental condition). CPM=counts per million reads, TF=transcription factor, TFBS=transcription factor binding site.

We next determined accessible conserved TFBS and corresponding expression of the TFs in microglia of (LPS-treated) Ercc1 deficient and WT mice. Compared to controls, SMAD1/3/4 binding sites are lost in microglia of KO mice (Figure 4C, Supplemental file 8), which are involved in maintenance of the microglia homeostatic gene signature (Butovsky et al., 2014; Massagué et al., 2005). Generally, immune activation of microglia results in the loss of the homeostatic signature (Holtman et al., 2015; Keren-Shaul et al., 2017; Krasemann et al., 2017; Zrzavy et al., 2017), and our data show that this is also true in primed microglia (Figure 4E). Microglial TF motifs with increased chromatin accessibility upon Ercc1 deletion include TFs whose associated functions were previously attributed to primed microglia (Holtman et al., 2015; D. D. A. Raj et al., 2014), namely lysosomal biogenesis (TFE3, Brady et al., 2018), inflammation (CEBP TF family (Simpson-Abelson et al., 2017; Straccia et al., 2011; Valente et al., 2013), IER TF family, ATF1) and proliferation (CEBP TF family, Gómez-Nicola et al., 2013; Pal et al., 2009) (Figure 4C, Supplemental file 8).

In contrast to microglia of LPS-treated WT mice, trained microglia of LPS-treated KO mice are enriched in accessible TF motifs for regulators with known roles in acute LPS-induced inflammation (Escoubet-Lozach et al., 2011; Holtman et al., 2015; Roach et al., 2007), including NFKB2 and REL/RELB, several members of the IRF TF family (IRF7, 8, 9), and IER-related TFs. In addition, ZEB1, associated with immune response suppression (C. J. Block et al., 2019; Wang et al., 2009), is depleted in trained microglia (Figure 4D, Supplemental file 8). Together with the fact that homeostatic genes in microglia of LPS-treated KO mice are even further downregulated than in KO microglia (Figure 4E), these results underline the training of microglia from KO mice.

These data suggest that Ercc1 depletion shapes a chromatin landscape that enables both the loss of the microglia homeostatic signature, and the gain of a transcriptional profile associated with inflammation, which is enhanced with LPS challenge.

### A large proportion of tolerized genes show an increased transcriptional response in primed and trained microglia

Both in tolerized (cluster 2 and 4, Figure 2A) and primed/trained gene sets (cluster 1, 2 and 4, Figure 2B), immune system processes were significantly enriched (Figure 2C, 2D). We intersected these gene sets and not only were similar biological processes affected, but many of the differentially regulated genes were also shared.

Out of the 302 tolerized genes, 145 genes overlap with acute LPS response-induced genes and 46 showed a significantly higher expression level in microglia of LPS-treated mice and *Ercc1^Δ/ko^* mice after LPS treatment. 264 and 249 genes were uniquely enriched in primed (KO-PBS versus WT-PBS) and trained (KO-LPS versus WT-LPS) microglia, respectively, and 251 genes overlapped between these conditions. Finally, 103 overlapping genes were enriched in acutely challenged, tolerized, primed as well as trained microglia (Figure 5A, Supplemental file 9). Significantly associated biological processes within these gene sets were identified (Figure 5B, Supplemental file 10). Genes involved in ‘organelle fission’, ‘nuclear division’ and ‘chromosome segregation’ were associated with and limited to primed microglia from Ercc1 deficient mice. The genes exclusive for training are involved in ‘response to oxidative stress’ and ‘ribosomal small subunit assembly’. The 251 genes that are shared between primed and trained microglia are associated with ‘positive regulation of cytokine production’ and ‘regulation of immune effector process’. The acute, tolerized and trained gene sets, with or without the primed gene set, share GO terms such as ‘regulation of innate immune response’, ‘NF-kappaB signaling’ and ‘regulation of apoptotic signaling pathway’.

**Figure 5.**
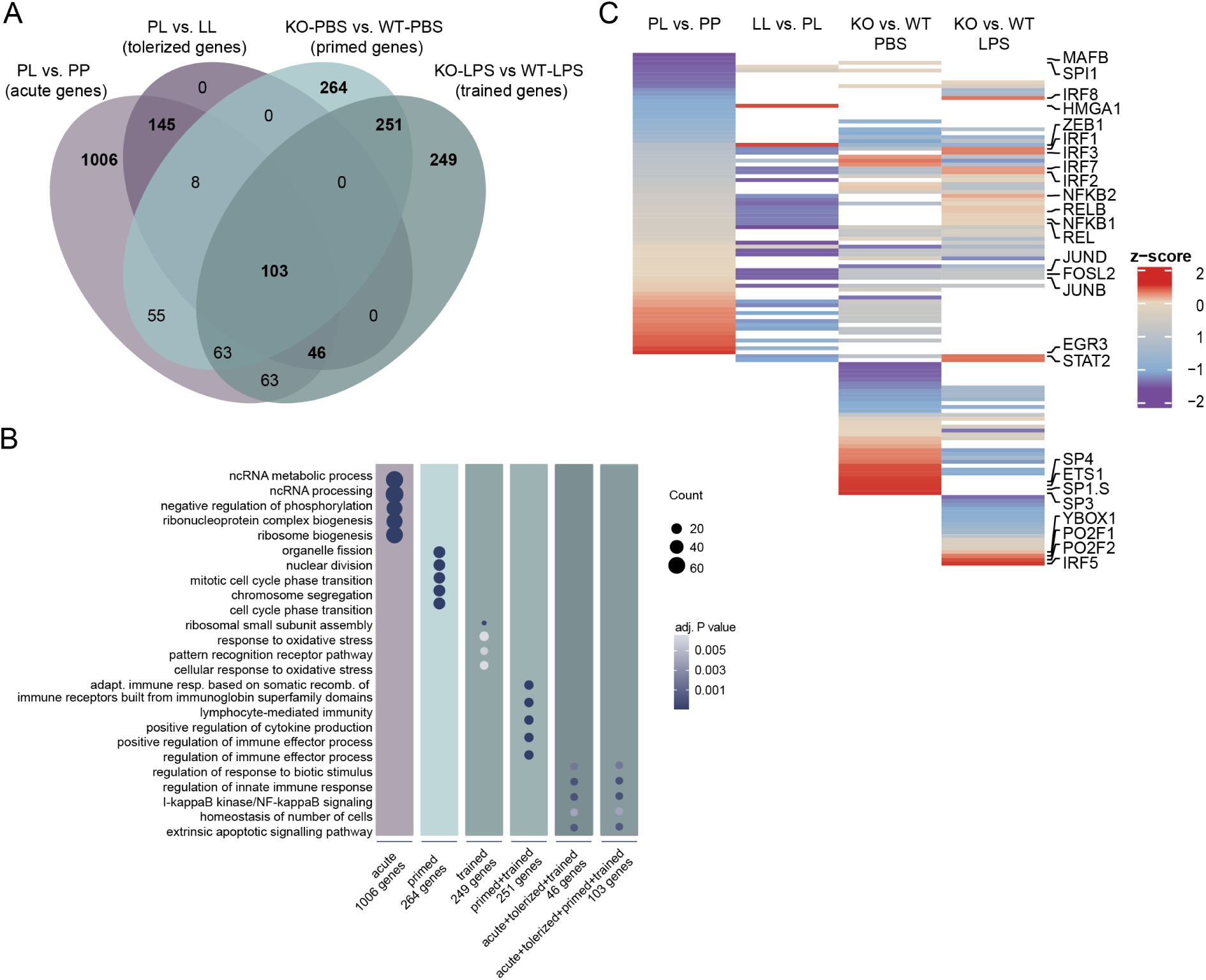
Inflammatory genes show distinct epigenetic regulation in ‘acute’, ‘tolerant’, ‘primed’ and ‘trained’ microglia. **a,** Venn diagram of the enriched genes in acute (PL versus PP, light purple, Supplemental file 1), tolerized (clusters 2 and 4, dark purple, Supplemental file 3), primed (KO-PBS versus WT-PBS, mint, Supplemental file 2) and trained (KO-LPS versus WT-LPS, dark mint, Supplemental file 2) microglial response. **b,** Dotplot depicting the GO terms associated with unique and overlapping gene sets of acute, tolerant, primed and trained microglia. The size of the dot represents the gene count per GO term and the color indicates the adjusted P-value. **c**, Heat map depicting row z-scores of weighted mean differences (adjusted P value<0.001) of ATAC peaks at loci of specific TF motifs in the indicated comparisons identified with diffTF (based on Figures 3B, 3D, 4C, 4D, Supplemental file 6, 8). Only accessible TF motifs of activating, repressing and undetermined TF are displayed (not-expressed TFs were filtered out). White space indicates non-significant weighted mean differences of ATAC peaks at the given locus and/or not-expressed TFs in the indicated comparison.

In order to determine possible regulators of the opposing LPS response between tolerized and trained genes, motifs for TFBS in genomic regions with enriched chromatin accessibility were identified with diffTF in acute (PL versus PP), tolerant (LL versus PL), primed (KO-PBS versus WT-PBS) and trained (KO-LPS versus WT-LPS) microglia (Figure 5C, Supplemental file 6, 8). The four identified microglial phenotypes (acute, tolerized, primed, trained) seem to be regulated by specific TF networks, explaining why gene sets, although being partially shared between some or all of the four phenotypes, are regulated in opposite directions.

Summarizing, the presented data indicate that microglia *in vivo* possess innate immune memory and that different types of stimuli, in this case Ercc1 deficiency or LPS, can leave epigenetic imprints which seem to influence the response towards a secondary challenge leading to microglia training or tolerance to LPS. Condition-specific epigenetic profiles seem to involve the activity of specific TF networks, which might drive the opposite regulation of shared genes in trained and tolerant microglia.

## Discussion

Monocytes and tissue resident macrophages play important roles in development, metabolism and immunity, thereby contributing to the maintenance of homeostasis. Though they are innate immune cells, macrophages can retain information of past inflammatory events, resulting in an altered response to reinfection. Depending on the primary trigger, macrophages can become ‘tolerant’, showing hypo-responsiveness, or ‘trained’ with increased responsiveness to subsequent stimuli. Biologically, these mechanisms are generally thought to provide a survival advantage in case of trained immunity (Netea, 2013), while the refractory state of tolerant macrophages causes increased mortality (Biswas & Lopez-Collazo, 2009). However, these effects seem to be context-dependent and it was hypothesized that trained immunity might have deleterious consequences in autoimmune diseases (Arts, Joosten, et al., 2018), whereas tolerance can provide a protective mechanism limiting the toxic effects of prolonged inflammation (Seeley & Ghosh, 2017).

Monocytes/macrophages undergo functional programming after exposure to microbial components (Kleinnijenhuis et al., 2012; Novakovic et al., 2016) and the associated genome-wide epigenetic characteristics of innate immune memory have been described over the past years (Arts, Moorlag, et al., 2018; Glass & Natoli, 2016; Novakovic et al., 2016; Perkins et al., 2016; Saeed et al., 2014). These observations are thought to provide clues as to which pathways to target to reverse ‘tolerance’ or stimulate ‘training’ in a clinical setting.

The CNS parenchyma contains microglia, tissue resident macrophages that fulfill highly specialized functions extending far beyond their innate immunological functions (Eggen et al., 2013; Eggen et al., 2017). Besides their different job-description that is attuned to their CNS environment, in contrast to some other tissue-derived macrophages, microglia also have a relatively long lifespan (Askew et al., 2017; Eggen et al., 2017; Füger et al., 2017; Tay et al., 2017). Microglia longevity together with the long-lasting nature of epigenetic mechanisms can have drastic effects on brain functioning and cognition.

In microglia, altered functional outcomes reminiscent of ‘tolerance’ and ‘training’ have been described and these mechanisms might contribute to poor cognitive outcomes in sepsis patients (Semmler et al., 2013), the general aged population and neurodegeneration (Haley et al., 2017; Knuesel et al., 2014; Neher & Cunningham, 2019; Pardon, 2015; Perry & Holmes, 2014; Wendeln et al., 2018). Particularly, disease features in mouse AD and stroke models appear to be altered in animals where microglia were exposed to systemic inflammatory stimuli (Wendeln et al., 2018).

Here, we show that exposure of microglia to an inflammatory challenge (LPS) or an environment of accelerated aging *in vivo* results in substantial transcriptional and epigenetic changes that impact on their future ability to mount an inflammatory response. In particular, we found that approximately 103 genes are oppositely regulated when ‘desensitized’ or ‘primed’ microglia are exposed to i.p. injection of LPS and that these genes are involved in inflammatory and apoptotic processes.

In the control situation, promoter and cis-regulatory elements controlling these inflammatory genes are characterized by a certain degree of chromatin accessibility, as well as H3K4me3 and H3K27Ac enrichment. In agreement with increased transcription of inflammatory genes in microglia from mice treated with LPS, these chromatin parameters were increased during the acute response. In case of tolerance, abundance of these marks is decreased after LPS re-exposure, which, at least partially, explains compromised induction of gene expression after secondary LPS challenge. Possibly, there is a second layer of gene expression repression by inhibitory histone marks. Previous data suggests a role for the inhibitory histone marks H3K9me2/3 in this context (Perkins et al., 2016; Schaafsma et al., 2015). The TF RELB has a recruiting role for H3K9me2/3 at the *Il1β* locus after LPS challenge, which leads to transcriptional repression of *Il1β* in response to a secondary LPS challenge (El Gazzar et al., 2008; Schaafsma et al., 2015). We identified enriched accessible binding motifs for REL and RELB in PL versus LL microglia genome-wide, indicating that a primary LPS challenge might lead to recruitment of REL/RELB at regulatory elements of tolerized genes and might inhibit gene expression upon secondary LPS challenge through recruitment of H3K9me2/3. However, this hypothesis needs to be confirmed in future ChIP-sequencing experiments.

In case of priming, continuous exposure to an aging environment results in increased chromatin accessibility as well as H3K4me3 and H3K27Ac enrichment. These changes are associated with increased gene expression levels in *Ercc1^Δ/ko^* microglia as well as after an LPS exposure. SMAD binding elements are known to act collaboratively with PU.1 and other TFs to facilitate transcription of the homeostatic microglia signature (Holtman et al., 2017). In the accelerated aging model, chromatin signatures associated with active gene expression are less associated with SMAD binding elements. This is accompanied by a decrease in expression of homeostatic microglia signature genes in *Ercc1^Δ/ko^* microglia, especially following LPS treatment. Microglia priming in this model is caused by neuronal genotoxic stress, since only Ercc1 deficiency in neurons, but not astrocytes and microglia induced microglia priming (D. D. A. Raj et al., 2014; Zhang et al., 2020). While active marks on promoters and enhancers correlate with increased expression, the Polycomb regulated repressive mark H3K27me3 is lost in some genes whose expression is increased in *Ercc1^Δ/ko^* microglia. Loss of the Polycomb mark H3K27me3 could be a critical determinant of cellular identity and function of primed microglia as the Polycomb repressive complex 2 (PRC2) is involved in maintenance of homeostatic microglia identity in different CNS brain regions. Loss of PRC2 activity in microglia resulted in aberrant gene expression and altered functionality (Ayata et al., 2018).

Besides the here presented data, microglia training was also observed in an AD amyloid mouse model, where an LPS challenge administered prior to the onset of AD pathology caused exacerbation of β-amyloidosis (Wendeln et al., 2018). Although the hyperresponsive nature of microglia to two stimuli seems to be comparable in these two studies, the underlying molecular mechanisms might be different due to the fact that the LPS stimulus and AD pathology were separated by a non-inflammatory phase (Wendeln et al., 2018), while persistent microglial activation is present in *Ercc1^Δ/ko^* mice.

Though the genes involved in tolerance and training are overlapping, the fact that the chromatin composition in these regions is diverse, suggests the involvement of distinct protein complexes and epigenetic enzymes. Summarizing, different molecular pathways and different epigenetic mechanisms regulate the behavior of inflammatory genes in ‘tolerant’ or ‘trained’ microglia.

Our data provides evidence that at least one type of macrophage, the CNS endogenous microglia, *in vivo* can adopt epigenetic programs that contribute to the establishment of different functional phenotypes and thereby influence neuroinflammation in the long-term.

## Supplementary figure legends

**Figure S1 related to figure 1.**
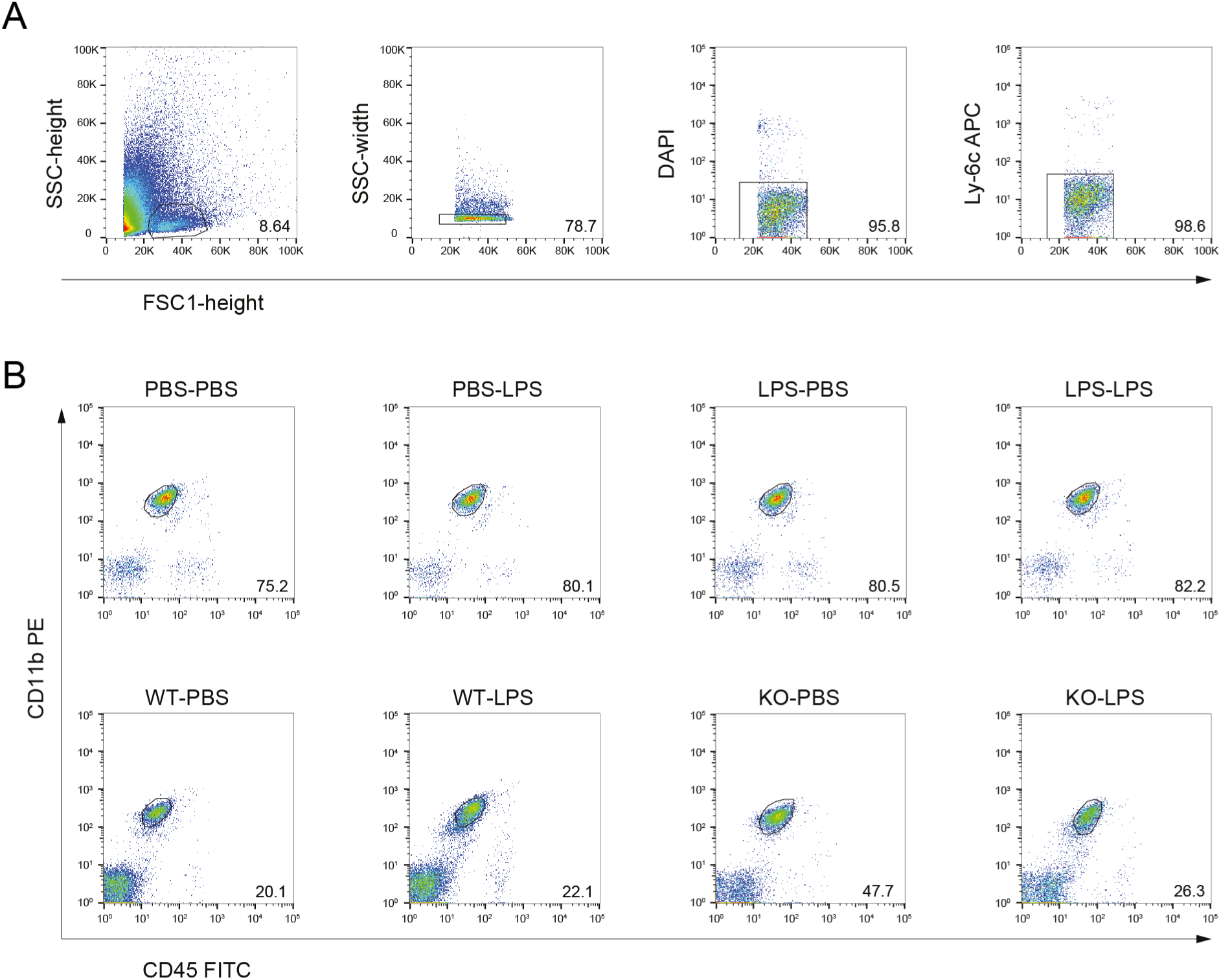
FACS sorting of microglia. **a,** Single, viable microglia are isolated using side scatter and forward scatter parameters, followed by exclusion of DAPI^pos^ (dead) events. Further purification was done by exclusion of Ly-6C^pos^ CNS macrophages. **b**, CD11b^pos^ and CD45^int^ microglia were sorted.

**Figure S2 related to figure 1.**
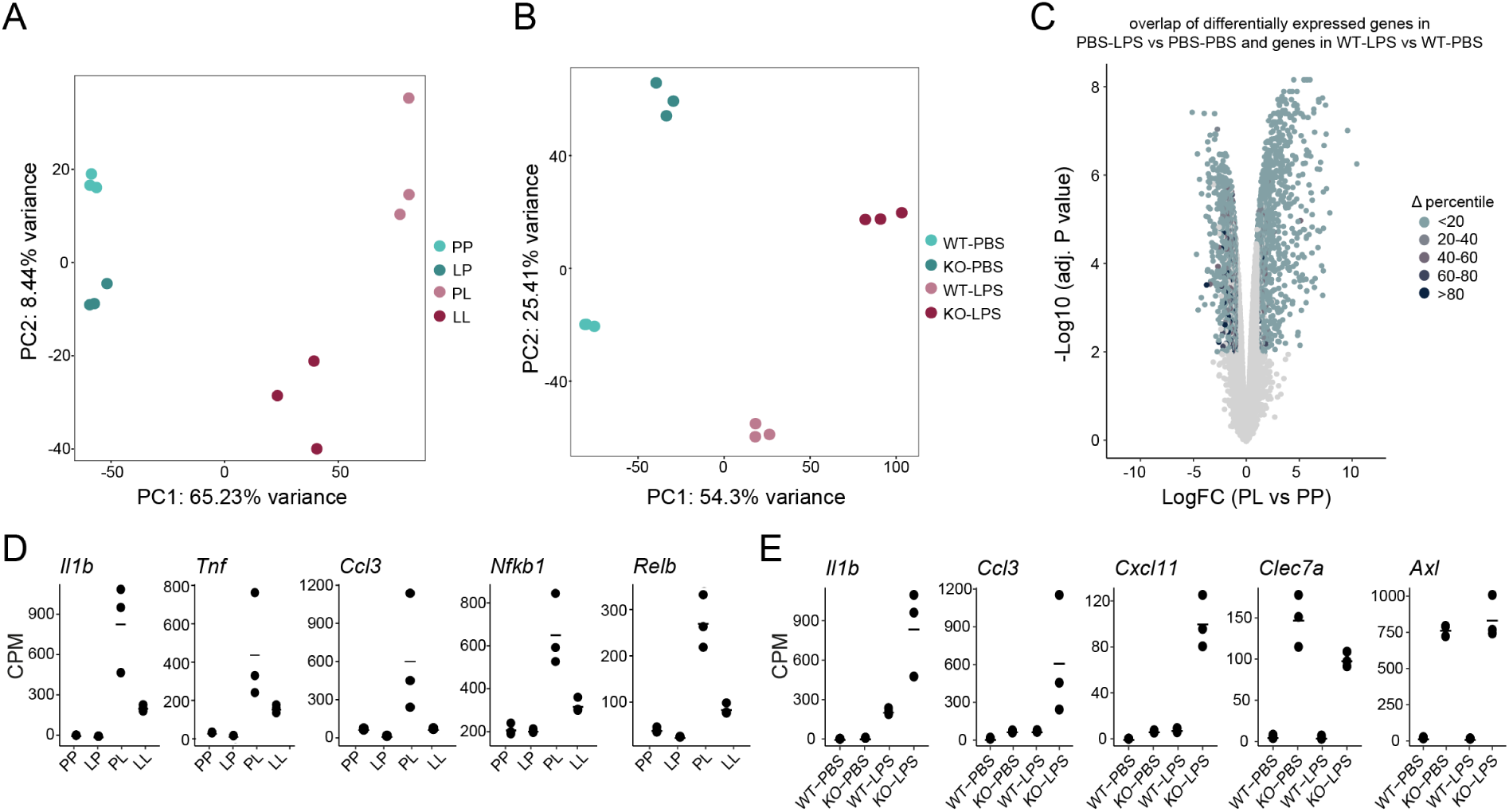
RNA-sequencing of desensitized/tolerant and primed/trained microglia. **a, b,** PCA-plots of RNA-seq data of microglia in the LPS desensitization tolerance (a) and Ercc1-induced priming (**b**) mouse models. Every dot depicts an individual animal (n=3 per experimental condition). **c,** Volcano plots illustrating the similarity in the acute LPS response in microglia from naive mice. Dots represent log fold change (LogFC) of differential expressed genes between PL and PP (**c**). Genes were ranked according to their expression level and based on that classified into percentiles. Next, for each gene of the two comparisons, the delta percentile was calculated and indicated as colors in the volcano plot, where light blue indicates similar and dark blue indicates deviant expression between the indicated conditions. Gray dots indicate gene expression differences with logFC<1 and adjusted P values>0.01. **d, e,** Gene expression values (CPM) of genes in the tolerance (**d**) or priming (**e**) mouse model. Every dot depicts an individual animal (n=3 per experimental condition). CPM= counts per million reads

**Figure S3 related to figure 2.**
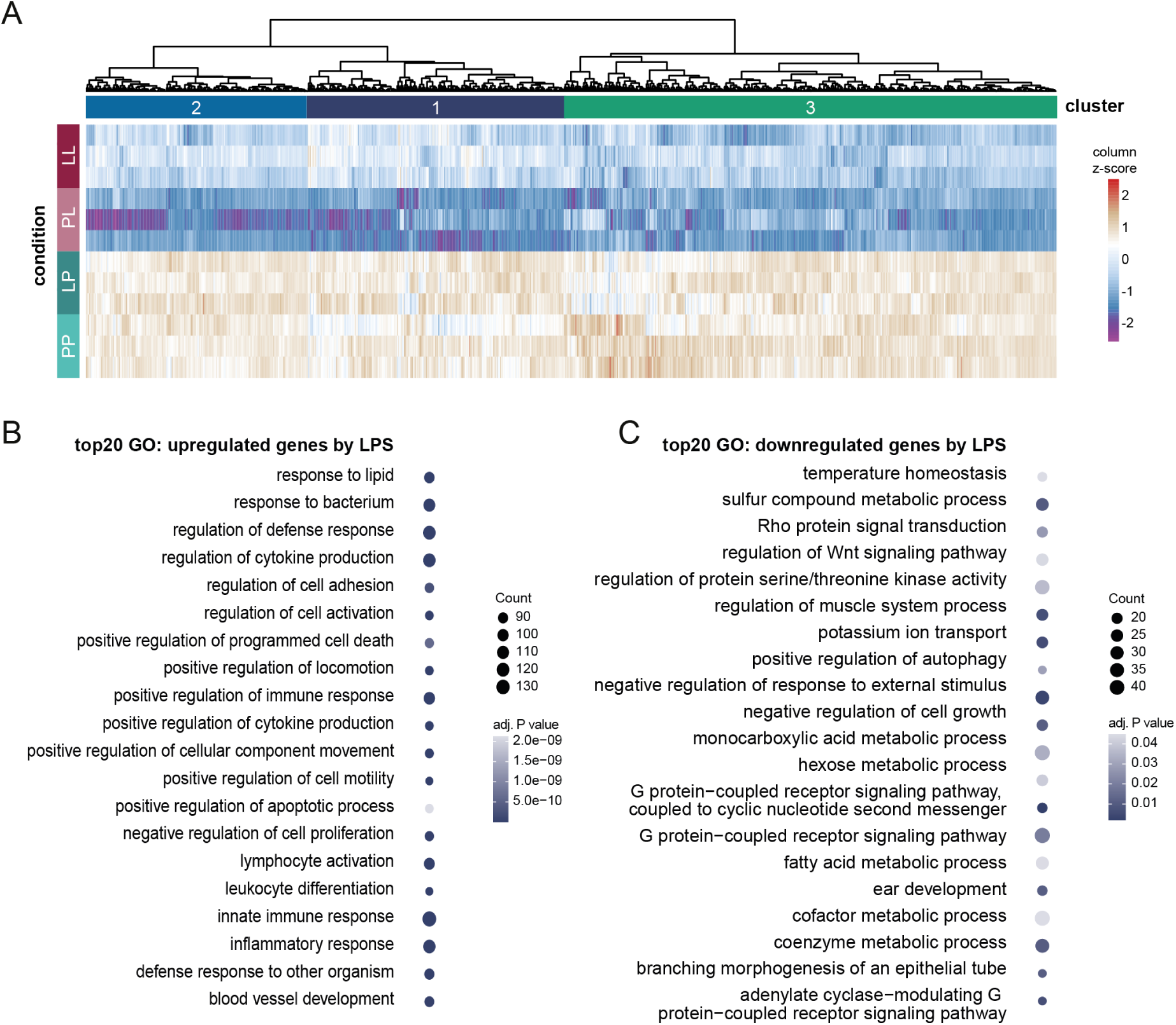
LPS-downregulated genes and associated GO terms. **a,** Heatmap with Manhattan distance-based hierarchical clustering analysis of downregulated genes in response to LPS in microglia of C57BL/6 mice (n=3 per experimental condition) three hours after i.p. injection with LPS (LogFC>1 and FDR<0.01, PP versus PL). **b, c,** Gene ontology (GO) analysis of genes upregulated (**b**) and downregulated (**c**) 3 hours after LPS challenge in microglia of C57BL/6 mice. Based on gene count per GO term, the top 20 GO terms were identified. The size of the dot represents the gene count per GO term and the color indicates the adjusted P value.

**Figure S4 related to figure 3 and 4.**
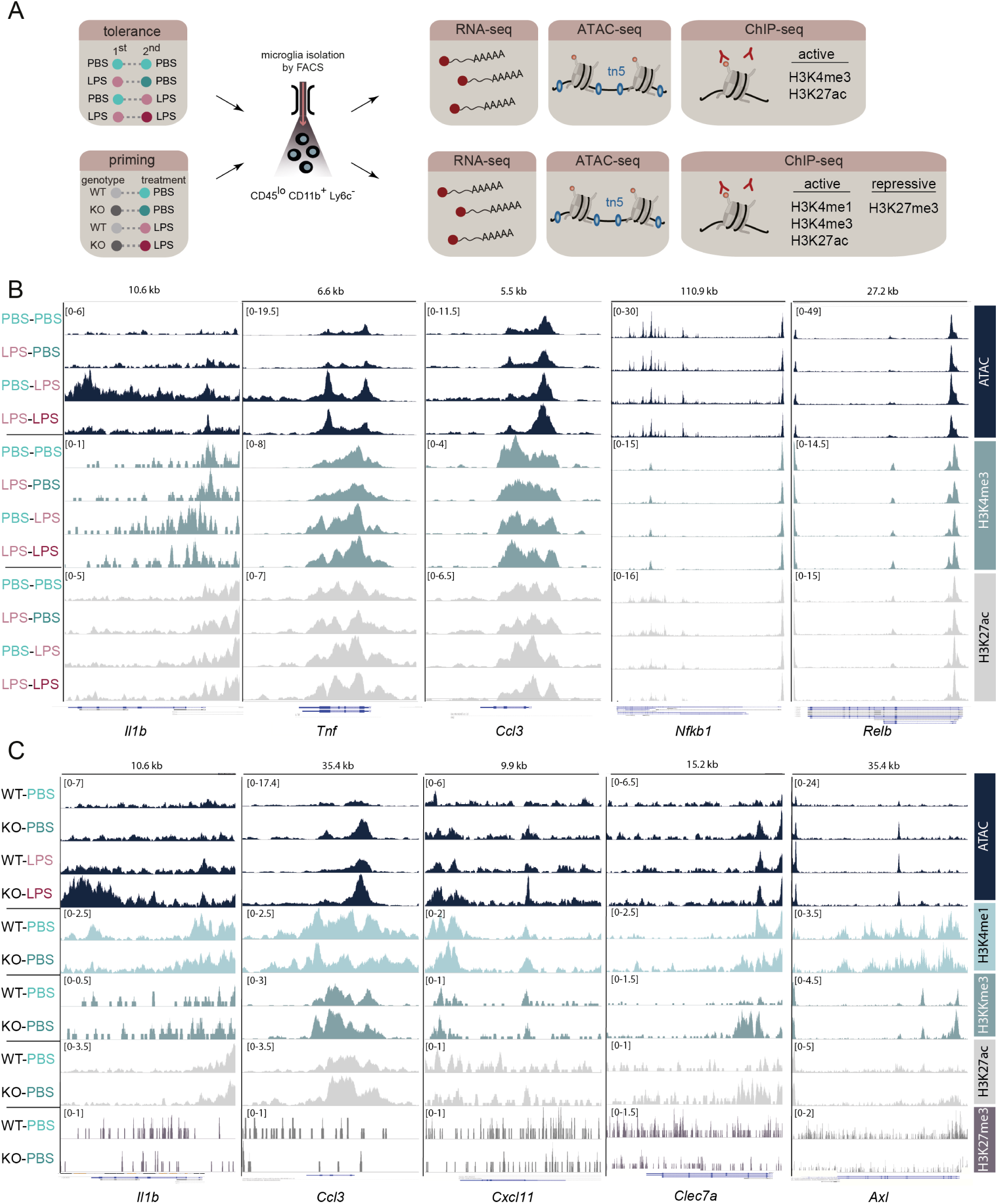
ATAC- and ChIP-sequencing peak enrichment at representative desensitized and primed gene loci. **a,** Experimental strategy for the analysis of chromatin accessibility, and occupation by histone modifications. H3K4me3 and H3K27ac were analyzed in ‘desensitized’ and ‘tolerant’ microglia. H3K4me1, H3K4me3 and H3K27ac were determined in ‘primed’ microglia. **b, c**, Tracks of ATAC and indicated histone marks sequencing data of representative desensitized/tolerant (**b**) and primed/trained (**c**) genes. For ChIP, chromatin of 5 mice per experimental group was pooled; for ATAC, microglia (80,000 total) from 2 mice per experimental group were pooled. Tracks were visualized using JetBrains SPAN peak analyzer. Gene expression of these genes are shown in S2D and S2E.

## Supplementary files

**Supplementary file 1**: count table and differentially expressed genes for all comparisons in the tolerance mouse model (related to Figure 1 and 3).

**Supplementary file 2**: count table and differentially expressed genes for all comparisons in the primed mouse model (related to Figure 1 and 4).

**Supplementary file 3:** Genes and associated gene ontology terms for each cluster identified in the tolerance model (related to figures 2 and S3).

**Supplementary file 4:** Genes and associated gene ontology terms for each cluster identified in the priming mouse model (related to figure 2 and S3).

**Supplemental file 5:** Annotated differential ATAC, H3K4me3 and H3K27ac peaks in microglia of PL versus PP, PL versus LL and LL versus LP mice (related to figure 3A, 3B, 3C).

**Supplemental file 6:** TFBS of differential ATAC peaks in microglia of PP versus PL, LL versus PL and LP versus LL mice and classification of TFs based on correlation of TFBS peaks with TF target gene expression (related to figure 3D, 3E, 3F, 5C).

**Supplemental file 7:** Annotated differential ATAC, H3K4me1, H3K4me3, H3K27ac and H3K27ac peaks in microglia of KO-PBS versus WT-PBS, KO-LPS versus WT-LPS, WT-LPS versus WT-PBS and KO-LPS versus WT-PBS mice (related to figure 4A, 4B).

**Supplemental file 8:** TFBS of differential ATAC peaks in microglia of WT-PBS versus KO-PBS and WT-LPS versus KO-LPS mice and classification of TFs based on correlation of TFBS peaks with TF target gene expression (related to Figure 4D, 4E, 5C).

**Supplemental file 9:** Lists of genes uniquely or overlappingly enriched in acute, tolerized, primed and trained microglia (related to figure 5A).

**Supplemental file 10:** GO terms associated with genes uniquely or overlappingly enriched in acute, tolerized, primed and trained microglia (related to figure 5B).

## Methods

### Animals

All the animal work was performed in the Central Animal Facility of the UMCG (CDP, Groningen, the Netherlands) and all animal-related studies were reviewed and approved by the Animal Care and Use Committee of the University of Groningen. Animals were conventionally housed under a 12/12 h light/dark cycle (8 p.m. lights off, 8 a.m. lights on) with *ad libitum* access to food and water.

#### Tolerance induction

Male C57BL/6J mice were obtained at the age of 7-9 weeks with weights in the range of 25-30 grams (Envigo, Horst, the Netherlands). Upon arrival, a minimum acclimatization time of 2 weeks was ensured, where mice were monitored weekly. All animals were housed individually and randomly assigned to experimental conditions.

To induce endotoxin tolerance, mice received 1 mg/kg body weight LPS (Sigma-Aldrich, *E. coli* 0111:B4, L4391) diluted in dPBS (Lonza, BE17512F) to a total volume of 200 µL by intraperitoneal injection. Immediately following LPS administration, mice were housed in a recovery cabinet at 26 °C for 24 hours. The weight and general health of injected animals were monitored daily until the body weight was completely restored (usually within 7 days). All control mice received 200 µL dPBS by intraperitoneal injection. After 4 weeks, the mice received a second injection with either dPBS or LPS (1mg/kg body weight, diluted in 200 µL dPBS).

#### Obtaining primed microglia

*Ercc1* transgenic mice (Weeda et al., 1997) were bred in house by crossing *Ercc1^wt/*292^* mice (FVB background, the *292 allele is hereafter indicated with Δ) with *Ercc1^wt/ko^* mice (BL6 background) as previously described (D. D. A. Raj et al., 2014). The offspring were genotyped after weaning using the primers listed in table 1. *Ercc1^Δ/ko^* were used as experimental mice while littermates with *Ercc1^wt/Δ^*, *Ercc1^wt/ko^* or *Ercc1^wt/wt^* genotypes were used as control. All the mice were group-housed in conventional cages. Initially, mice were monitored weekly, which increased to twice per week after the aging-related symptoms appeared. Bottles with long drinking spouts were provided to prevent dehydration of *Ercc1^Δ/ko^* animals. At 11-12 weeks of age, the mice received 1 mg/kg body weight LPS or dPBS as described above. Immediately following LPS administration, mice were temporarily housed in a recovery cabinet at 26 °C.

**Table 1.**
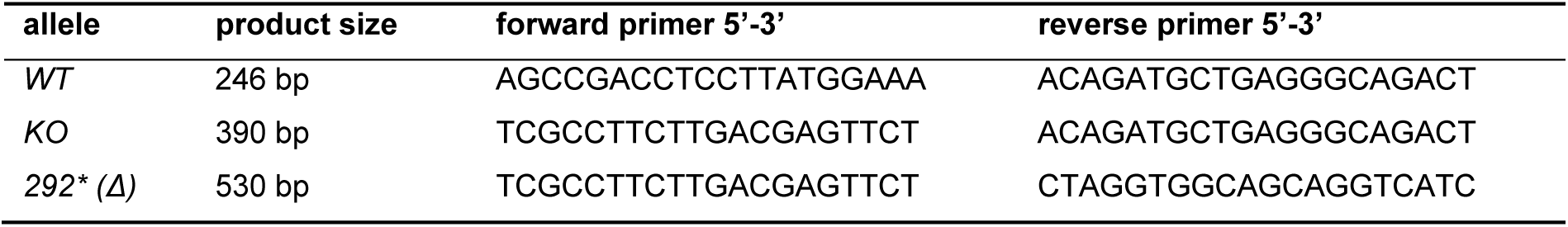
*Ercc1* genotyping primers.

All animals were sacrificed under deep anesthesia (4% isoflurane with 7.5% O_2_) and perfused with cold dPBS exactly three hours after the last injection.

### Microglia isolation and flow cytometry

Microglia were isolated as previously described (Galatro, Vainchtein, et al., 2017). After perfusion, brains were removed from the skull and kept in cold medium A (HBSS (Gibco, 14170-088) with 0.6% glucose (Sigma, G8769) and 7.5 mM HEPES (Lonza, BE17-737E)). All subsequent steps were performed on ice, centrifugation was at 4°C. Brains were dissociated using a potter-elvehjem tissue homogenizer after which the homogenate was passed over a 70 µM cell strainer (Corning, 352350) and pelleted by centrifugation at 220 xg for 10 min. Next, myelin was removed by resuspending the pellet in 25 mL 24% percoll (Fisher, 17-0891-01) in medium A (1x final concentration) with 3 mL PBS layered on top, followed by centrifugation for 20 min at 950 xg (acceleration 4 and brake 0). The microglia enriched cell pellets were incubated with CD11b-PE (clone M1/70, eBiosciences, 12-0112-82), CD45-FITC (clone 30-F11, eBiosciences, 11-0451-82), and Ly6c-APC (clone HK1.4, Biolegend, 128016) antibodies for 30 min on ice. Then the cells were washed once in medium A without phenol red and filtered into FACS tubes. Microglia were sorted by gating the DAPI^neg^CD11b^high^CD45^int^Ly6c^neg^ cells using the Beckman Coulter MoFlo Astrios or XDP. Microglia were collected in siliconized Eppendorf tubes (Sigma, T3406-250EA) containing medium A. Flow cytometry data was analyzed using FlowJo Analysis Software.

### RNA isolation and RNA sequencing

Total RNA was isolated using a Qiagen RNeasy Micro Kit (Qiagen, 74004) according to the manufacturer’s instructions. cDNA was synthesized using random hexamer primers (Thermo, SO142), dNTPs (Thermo, R0192) and M-MuLV Reverse Transcriptase (Thermo, EP0442) in the presence of Ribolock RNase inhibitor (Thermo, EO0382). Quantitative PCR reactions were performed using iTaq mastermix (Biorad, 172-5125) on ABI7900RH or QuantStudio 7 (Applied Biosystems) or LightCycler^®^ 480 (Roche) PCR systems.

#### Endotoxin tolerance

The quality of the total RNA was determined using an Experion (Biorad). All included samples had an RNA quality indicator > 6. Sequencing libraries were generated with a TruSeq RNA library prep kit (Illumina, RS-122-2001). Pooled libraries were sequenced with a HiSeq Rapid SBS kit (50 cycles, Illumina, FC-402-4022) using single reads on a HiSeq 2500 (Illumina).

#### Priming/Ercc1 knockout

The quality of total RNA samples isolated from *Ercc1^Δ/ko^* mice was determined on a LabchipGX (PerkinElmer). All included samples had an RNA quality score > 5. Sequencing libraries were generated using NEXTflex^®^ Rapid Illumina Directional RNA-Seq Library Prep Kit (BiooScientific, NOVA-5138-10) with polyA selection. Pooled libraries were sequenced using the NextSeq 500/550 High Output v2 kit (75 cycles, Illumina, FC-404-2005) with single reads on a NextSeq500 (Illumina).

### RNA-sequencing analysis

Samples (n=3 per condition) were processed using our *in-house* pipeline, where quality control was performed with FastQC (v0.11.8). Adapter sequences and were removed with bbduk (v38.69). The Ensembl genome *Mus musculus* (GRCm38.82) was used for alignment (STAR v2.7.3a, Dobin et al., 2013). Sorting of the aligned reads was done with bamsort tool from biobambam2 tools (v2.0.95). featureCounts (v2.0.0, Liao et al., 2014) was used to quantify the reads. Picard (v1.130, “Picard Toolkit.,” 2019) and samtools were used to perform the quality control check and the generation of the fastq files. Downstream analyses were performed using R/Bioconductor packages (v.3.11), as briefly summarized. Specific functions from EdgeR (v3.30.3, Robinson et al., 2009) were used for data normalization and calculation of rpkm expression values. Log2-CPM values and mean-variance relationship were calculated with the voomWithQualityWeights function from limma (v.3.44.3, Ritchie et al., 2015). Unwanted/hidden sources of variation were removed using sva (v3.36.0, Leek et al., 2020). Differential expression analysis was performed using limma. Annotation was performed with biomaRt (Durinck et al., 2009). For plotting purposes, genes with a logFC>1 and FDR<0.01 were considered differentially expressed.

Clustering analysis were performed using the ward.D2 clustering method and Manhattan distance as clustering metrics. Heat maps were assembled using the pheatmap package (v1.0.12, Kolde, 2019). The optimal number of gene clusters was estimated upon visual inspection of the heat maps. Gene ontology enrichment analysis of the gene clusters was performed with the ‘enrichGO’ function of the clusterProfiler package (v3.16.1, Yu et al., 2012). PCA plots, scatterplots and dotplots were made with the package ggplot2 (v3.3.2, Wickham et al., 2016) and standard plot functions from R.

To visualize the overlap of differentially expressed genes of different comparisons (Figure S2C), gene lists of the indicated comparisons were ranked based on expression level. Following, percentiles were assigned to the ranked genes and Δpercentiles were calculated by subtracting percentiles of each gene from the two indicated comparisons. The results were depicted in a volcano plot, where the dots are differential expressed genes (logF>1, FDR<0.01) in the indicated comparison and the color of the dots shows overlap of gene expression levels from the two indicated comparisons.

For the Venn diagram in Figure 5a, only upregulated genes of the PL versus PP, KO-PBS versus WT-PBS and KO-LPs versus WT-LPS differential gene lists (Supplementary file 1 and 2) and genes of cluster 2 and 4 of the tolerance moue model (Supplementary file 3) were use. The Venn diagram was made with the ‘venn’ function of the gplots package (v3.1.0, Warnes et al., 2015).

### ChIP-sequencing

The procedure of Chromatin immunoprecipitation has been described previously (Schaafsma et al., 2015). Sorted microglia were fixed in 1 mL 1% formaldehyde diluted in dPBS at 20°C for 10 min and fixation was stopped by adding glycine to a final concentration of 0.125 M glycine. Fixed cells were washed twice by 1 mL cold dPBS, and then lysed with cell lysis buffer (5 mM Pipes, pH8.0; 85 mM KCl; 0.5% NP-40) by incubating on ice for 10 min. At the end, the cells were lysed in 250 µL nuclear lysis buffer (50 mM Tris.HCl, pH8.1; 10 mM EDTA, pH8.0; 1% SDS) to obtain the crosslinked chromatin. Chromatin was sonicated using a Bioruptor (Diagenode) at “high” power for 20 min (30 sec on and 30 sec off, for 20 cycles) at 4°C. Chromatin from animals within the same treatment group was pooled (5 animals per pool) and precleared using protein A agarose beads (25%, diluted in ChIP dilution buffer; Protein A Agarose/Salmon Sperm DNA, Millipore, 16-157) and incubated overnight at 4°C (final buffer composition during antibody incubation was 0.1% SDS; 1% Triton-X-100; 2.4 mM EDTA; 20 mMTris.HCl, pH8.1; 150 mM NaCl) with antibodies for specific histone modifications (the information of antibodies is available in table 2, the specificity of antibodies have been checked for H3K27me3 peptides, the information of these peptides is listed in table 2). The chromatin incubated with IgG was used as negative control while the chromatin saved without antibody incubation served as input. The next day, immune complexes were precipitated with 80 µL protein A beads (25%) for 2 h at 4 °C, washed by low salt wash buffer (150 mM NaCl; 0.1% SDS; 1% Triton-x-100; 2 mM EDTA, pH8.0; 20 mM Tris.HCl, pH8.1), high salt wash buffer (500 mM NaCl; 0.1% SDS; 1% Triton-x-100; 2 mM EDTA, pH8.0; 20 mM Tris.HCl, pH8.1), LiCl wash buffer (0.25 M LiCl; 1% NP-40; 1% Na-deoxycholate; 1 mM EDTA; 10 mM Tris.HCl, pH8.1), and TE (10 mM Tris.HCl, pH8.0; 1 mM EDTA, pH8.0). After the chromatin was eluted from the beads, the precipitated chromatin was de-crosslinked overnight at 65 °C. Afterwards, RNase A (Thermofisher, EN0531) and Proteinase K (Sigma, P2308) were added. Finally, the DNA was purified by GeneJET PCR purification kit (ThermoFisher, k0701).

**Table 2.** Antibodies used for ChIP.

| antibody | supplier | full name | cat # | lot # |
| --- | --- | --- | --- | --- |
| H3K4me1 | Abcam | Anti-Histone H3 (mono methyl K4) antibody – ChIP Grade | Ab8895 | GR193737-1 |
| H3K4me3 | Millipore | Anti-trimethyl-Histone H3 (Lys4) | 07-473 | 2117175 |
| H3K27ac | Abcam | Anti-Histone H3 (acetyl K27) antibody – ChIP Grade | Ab4729 | GR200563-1 |
| H3K27me3 | Millipore | ChIP Ab+™ Trimethyl-Histone H3 (Lys27) | 17-622 | 2325081 |

Sequencing libraries were generated from the purified DNA by MicroPlex Library Preparation Kit v1 x12 (Diagenode, C05010010) for tolerized samples or MicroPlex Library Preparation Kit v2 x12 (Diagenode, C05010012) in case of primed samples. The libraries were quantified by Agilent 2100 Bioanalyzer, pooled and sequenced with a HiSeq Rapid SBS kit (50 cycles, Illumina, FC-402-4022) using single reads on a HiSeq 2500 (Illumina).

### ATAC sequencing

ATAC-sequencing libraries were generated using Nextera^®^ DNA Sample Preparation Kit (Illumina, FC-121-1030) following the methods described by Buenrostro et al., 2013, 2015. A total number of 80,000 microglia were pooled from two animals (40,000 cells from each) and collected in Eppendorf tubes containing 300 µL medium A. Cells were pelleted by centrifugation (10 min, 4 °C, 500 xg), resuspended in 50 μL of cold lysis buffer (10 mM Tris-HCl, pH 7.4, 10 mM NaCl, 3 mM MgCl_2_, 0.1% IGEPAL CA-630) and immediately centrifuged as before. Next, nuclei were resuspended in 50 μL transposition reaction mix (1x TD reaction buffer, 2.5 μL TN5 transposase) and incubated at 37°C for 30 min. Immediately following transposition, the DNA was purified using a minElute PCR purification kit (Qiagen, 28004) following the manufacturer’s instructions. The transposed DNA fragments were further amplified and barcoded (Buenrostro et al., 2013, 2015) and purified with a ChIP DNA Clean & Concentrator kit (Zymo, D5205). The fragments were run on 2% E-Gel™ EX agarose gels (Thermo Fisher scientific, G521802) and 150-600 bp fragments were excised, followed by purification with Zymoclean™ Gel DNA Recovery Kit (Zymo, D4007). Library concentration was determined with an Agilent 2100 Bioanalyzer after which 8 samples were pooled and sequenced using HiSeq Rapid SBS Kit v2 (50 cycles) using paired end reads on a HiSeq2500 (Illumina).

### ChIP- and ATAC-sequencing analysis

ATAC and ChIP samples were aligned to the *Mus musculus* genome (mm10/GRCm38) with the use of Bowtie 2 (v2.3.5.1, Langmead & Salzberg, 2012) using the –very-sensitive flag. Bamsort and bammarkduplicates from biobambam2 tools (v2.0.95) were used to sort the aligned files and to remove duplicated reads. Samtools (v.1.1.0, Danecek et al., 2021) was used to remove low quality (q<30) and blacklisted alignments. For Chip-seq data, peak calling was performed using the JetBrains SPAN peak analyzer (v.0.11.0) using default parameters, which were later manually refined upon visual inspection using the JetBrains JBR browser (v.1.0 beta) on each sample. BigWig files were generated using deepTools bamCoverage (v.3.5.0) with RPGC normalization. ATAC-seq peaks were called using Genrich (v.0.5) with the ATAC-seq mode (-j switch), and -p parameter set to 0.01. Differential peak calling for ChIP- and ATAC-seq were performed with manorm (v.1.3.0, Shao et al., 2012). The annotation of differential peaks was performed with the annotatePeaks function from R/Bioconductor ChIPseeker package (v1.24.0, G. Yu et al., 2015).

Analysis of differential transcription factor binding sites accessibility and classification of transcription factors into activators, repressors or undetermined was performed with the diffTF package (v1.7.1., Berest et al., 2019) based on ATAC- and RNA-seq data.

ChIP- and ATAC-seq peaks are visualized with the JetBrains SPAN peak analyzer. The heatmap in Figure 5c is based on the diffTF output in Supplemental file 6 & 8. The row z-score was calculated from weighted mean differences of ATAC peaks from the indicated comparison. Following, in each comparison non-significant differential peaks (adjusted P value>0.001) and TF classified as ‘not-expressed’ were omitted. The row z-scores of significant differential accessible regions (adjusted P value>0.001) of putative transcriptional activators, repressors and undetermined TFs are visualized in a heatmap assembled with the ‘Heatmap’ function of the ComplexHeatmap package (v2.4.3, Gu et al., 2016).

## Acknowledgements

The authors thank Karina Hoekstra-Wakker, Nancy Halsema, and Diana Spierings for sequencing support, Geert Mesander, Henk Moes, and Roelof Jan van der Lei for technical assistance with FACS sorting, and Hilmar RJ van Weering for artwork. This work was supported by a China Scholarship Council fellowship to XZ (Grant # 201306300082) and a Graduate School of Medical Sciences-UMCG scholarship to LK. SMK is funded by the Netherlands Organization for Scientific Research (NWO, VENI, #016.161.072) and the MS Research Foundation (16-947).

## Author contributions

Conceptualization, X.Z., S.M.K., B.J.L.E.; Formal Analysis, A.M.L., M.L.D., L.K.; Investigation, X.Z., S.M.K., L.K., N.B., E.M.W.; Writing – Original Draft, S.M.K., B.J.L.E.; Writing – Review & Editing, X.Z., L.K., B.J.L.E., S.M.K., A.M.L., M.L.D., N.B., E.M.W., H.W.G.M.B.; Supervision, S.M.K., B.J.L.E.; Funding Acquisition, X.Z., S.M.K., H.W.G.M.B., B.J.L.E.

## Data availability statement

All next-generation sequencing data can be viewed at NCBI GEO under accession number GSE175578.

